# Hemodynamic and electromechanical effects of paraquat in rat heart

**DOI:** 10.1101/2020.06.01.127464

**Authors:** Chih-Chuan Lin, Kuang-Hung Hsu, Gwo-Jyh Chang

## Abstract

Paraquat (PQ) is a highly lethal herbicide. Ingestion of large quantities of PQ usually results in cardiovascular collapse and eventually death. However, the mechanism of acute PQ poisoning induced cardiotoxicity is poorly understood. Therefore, the purpose of the present study was to aim to investigate the mechanisms of PQ induced cardiotoxicity by examining the effects of PQ on hemodynamics in vivo, as well as in vitro on isolated hearts and ventricular myocytes in rats. Intravenous administration of PQ (100 or 180 mg/kg) in anesthetized rats induced dose-dependent decreases in heart rate, blood pressure, and cardiac contractility (left ventricular [LV] d*P*/d*t*_max_). Furthermore, it prolonged the rate-corrected QT (QTc) interval. In Langendorff-perfused isolated hearts, PQ (33 or 60 μM) decreased LV pressure and contractility (LV d*P*/d*t*_max_ in isolated ventricular myocytes), PQ (10–60 μM) decreased the amplitude of Ca^2+^ transients and cell shortening in a concentration-dependent manner. Patch-clamp experiments demonstrated that PQ decreased the amplitude and availability of the transient outward K^+^ channel (*I*_*to*_) and altered its gating kinetics. These results suggest that PQ-induced cardiotoxicity results mainly from diminished Ca^2+^ transients and inhibited K^+^ channels, which lead to the suppression of LV contractile force and prolongation of the QTc interval.

## Introduction

Paraquat (PQ)is a highly toxic herbicide causes significant mortality when ingested. It is an widely used herbicides because of its excellent ability to eliminate weeds [1]. However, intentional ingestion of PQ to commit suicide is a common problem in some Asian countries, such as Taiwan, Japan, South Korea, Malaysia, and Sri Lanka [2–5].

Fulminate PQ poisoning patients with cardiovascular collapse and multiple organ failure have the highest mortality rate. Death usually occurs within 24–72 h mostly or 1 week [6,7]. Histologically, acute PQ poisoning may cause PQ accumulation and direct myocardium injury [8–10]. Acute PQ poisoning also causes rate-corrected QT (QTc) interval prolongation and serves as a useful prognostic factor for poor outcome clinically [11]. QTc prolongation are thought to be related to left ventricular (LV) systolic dysfunction in cardiac disease patients, too [12]. In human PQ poisoning cases, form minimal changes on the ECG to extensive myocardial necrosis had been observed [13]. However, the pathophysiologic mechanism of PQ caused cardiac toxicity is less studied. Therefore, the aim of this study was to investigate the mechanisms of PQ induced cardiotoxicity by examining the hemodynamic effects of PQ in vivo, as well as the electromechanical effects on isolated rat hearts and ventricular myocytes in vitro. Several approaches were used to achieve the aims. First, the effects of PQ on hemodynamic parameters and electrocardiogram (ECG) will be demonstrated in anesthetized rats. then, the PQ’s effects on LV pressure of isolated Langendorff-perfused hearts, as well as its effects on Ca^2+^ transients, cell shortening, and K^+^ currents of ventricular myocytes will also be investigated.

## Materials and Methods

All experiments were approved by the Institutional Animal Care and Use Committee of Chang Gung University (Tao-Yuan, Taiwan) and performed in accordance with the Guide for the Care and Use of Laboratory Animals published by the US National Institutes of Health (NIH publication No. 85-23, revised 1996). Animals were housed in polypropylene cages with a controlled light and dark cycle of 12 hours each at 24–26°C. Food and water were available ad libitum. A total of 93 animals were used in the experiments described here.

For this study, Adult male Sprague Dawley rats weighing 250–300 g purchased from Labsco Tech. Co. (Taipei, Taiwan) were anesthetized with urethane (1.25 g/kg, i.p.).

### Materials

PQ was purchased from Sigma-Aldrich (St. Louis, MO, USA) and prepared in physiological saline. Fura-2-acetoxymethyl ester (Fura-2-AM) and Pluronic F-127 were purchased from Molecular Probes (Eugene, OR, USA) and dissolved in dimethylsulfoxide (DMSO). All other drugs were purchased from Sigma-Aldrich and prepared in physiological saline before the start of each experiment. The LD_50_ value in rats was reported to be around 57 mg/kg. Therefore, we used 100 and 180 mg/kg (approximately twice and triple LD_50,_ respectively) for the in vivo study. Accordingly, dose ranges in vitro for the PQ at doses of study were also calculated.

### Hemodynamic and ECG measurements in anesthetized rats

Adult male Sprague Dawley rats weighing 250–300 g purchased from Labsco Tech. Co. (Taipei, Taiwan) were anesthetized with urethane (1.25 g/kg, i.p.). A polyethylene (PE 50) cannula filled with heparinized saline (25 IU/mL) was inserted into the femoral artery to measure arterial pressure. The arterial cannula was connected to an MLT0380/D pressure transducer (ADInstruments, Bella Vista, Australia) linked to a QuadBridge amplifier (ADInstruments). ECG needles were connected to a biological amplifier (ADInstruments) and lead II ECGs were recorded simultaneously. QT intervals were rate corrected according to Bazett’s formula where QTc = QT/√RR. For measurement of LVP, a 1.9-F microtip pressure–volume catheter (Model FTS-1912B, Scisense Inc., London, Canada) was advanced from the right carotid artery into the LV under pressure control. LP pressure (LVP) signals were continuously recorded using a pressure–volume conductance system (Model 891A, Scisense). Output signals from these amplifiers were connected to a Ponemah ACQ16 acquisition system (DSI Ponemah, Valley View, OH, USA) and recorded at a sampling rate of 4 kHz, and then stored and displayed on a computer. All arterial and LV pressure data were analyzed using a data analysis program (P3 Plus4.80-SP4, DSI Ponemah) and the mean arterial blood pressure (MAP), heart rate (HR), systolic blood pressure (SBP), diastolic blood pressure (DBP), LV end-systolic pressure (LVESP), LV end-diastolic pressure (LVEDP), and maximal rate of rise (+d*P*/d*t*_max_) and fall (−d*P*/d*t*_max_) of LVP were computed. The femoral vein was cannulated with a PE50 catheter for drug administration. After 20 min of stabilization, a saline or a PQ (100 or 180 mg/kg) solution was administered intravenously in the control or treated groups, respectively. The volume of infusion was 1 mL/kg and given for 1 min.

### Intraventricular pressure measurement in Langendorff-perfused rat hearts

The rats was anesthetized with pentobarbital sodium (50 mg/kg, i.p.) and placed on an operating table. Rat hearts were excised immediately, mounted on a Langendorff apparatus, and perfused at a constant pressure of approximately 55 mmHg with oxygenated (95% O_2_ and 5% CO_2_) normal Tyrode solution containing (in μM): NaCl 137.0, KCl 5.4, MgCl_2_ 1.1, NaHCO_3_ 11.9, NaH_2_PO_4_ 0.33, CaCl_2_ 1.8, and dextrose 11.0 at 37°C as described previously[14]. A latex balloon (size No. 5, Radnoti, Monrovia, CA, USA) connected by a short stainless steel tube to a pressure transducer (P23XL-1, Becton, Dickinson & Co., Franklin Lakes, NJ, USA) was inserted into the LV cavity via the left atrium. The balloon was inflated with 0.04 mL distilled water, which was sufficient to produce an end-diastolic pressure of 8–12 mmHg. The ventricles were paced electrically at a rate of 300 beats per min by platinum contact electrodes positioned on the right ventricular apex. Data were recorded on a WindowGraf recorder (Gould Inc., Cleveland, Ohio, USA) and digitized with a computer-based data acquisition system (PowerLab/4SP with Chart 5 software, ADInstruments). Each preparation was allowed to equilibrate for 2–2.5 h before drug testing. LV developed pressure (LVDP) was calculated by subtracting LVEDP from the LV peak systolic pressure. Differentiation of the LVP signal was used to determine LV + d*P*/d*t*_max_ and −d*P*/d*t*_max_.

### Single cardiac myocyte isolation

Single ventricular myocytes from adult rats were obtained by an enzymatic dissociation method described previously [14]. In brief, the excised heart was mounted on a Langendorff apparatus and retrogradely perfused at a rate of 6 mL/(min·g cardiac tissue) by a peristaltic pump with nominally Ca^2+^-free HEPES-buffered Tyrode solution containing (in μM): NaCl 137.0, KCl 5.4, KH_2_PO_4_ 1.2, MgSO_4_ 1.22, dextrose 22.0, and HEPES 6.0, titrated to pH 7.4 with NaOH. The perfusate was oxygenated and maintained at 37 ± 0.2°C by a heating circulator. After 5 min, the perfusate was changed to the same solution containing 0.3 mg/mL collagenase (Type II, Sigma-Aldrich) and 0.1 mg/mL protease (Type XIV, Sigma-Aldrich). After digestion for 7–15 min, the residual enzyme solution was removed by perfusing 0.05 μM Ca^2+^-containing Tyrode solution. Thereafter, the ventricles were separated from the atria, dispersed, and stored in 0.2 μM Ca^2+^-containing Tyrode solution for later use. Rod-shaped Ca^2+^-tolerant viable cells with clear striations were used for experiments.

### Measurements of intracellular Ca^2+^ transients and cell shortening

Ventricular myocytes were loaded with the fluorescent Ca^2+^-sensitive indicator, fura-2, by incubating cells in 0.5 μM Ca^2+^-containing HEPES solution containing 5 μM fura-2-AM and 2% Pluronic F-127 for 30 min at room temperature, as described previously [15]. After washing out the excess fura-2-AM, cells were stored in 0.5 μM Ca^2+^-containing HEPES solution. Fura-2–loaded myocytes were transferred to 1.8 μM Ca^2+^-containing HEPES buffer for at least 30 min before beginning the experiments.

Myocytes were then electrically stimulated using a pair of platinum electrodes with a 2-ms and 2-fold threshold rectangular voltage pulse at 1 Hz. Myocytes were placed on an inverted microscope (Axio Observer Z1, Carl Zeiss MicroImaging GmbH, Jena, Germany) equipped with a heated (37°C) chamber. The cells were illuminated with ultraviolet light from a light source (DeltaScan, Photon Technology International [PTI], NJ, USA). The excitation lights with a wavelength of 340 or 380 nm passed through a ×40 oil immersion objective to the cell by a dichroic mirror. The emission light passed through a filter (510 nm) and was detected by a photomultiplier tube and recorded by using a RatioMaster fluorometer (PTI). To minimize photobleaching of fura-2, excitation light was applied intermittently and attenuated by 90% with the use of a neutral density filter. Signals were acquired using a data acquisition system controlled with professional software (FeliX32™, PTI). Since the fura-2 ratio is not a linear function of intracellular Ca^2+^ concentration when cells are loaded with fura-2-AM, intracellular Ca^2+^ was directly expressed as the ratio of the light emitted at excitation wavelengths 340 nm and 380 nm (*F*_340_/*F*_380_). Background fluorescence measured from a cell-free field was subtracted from all recordings before calculation of ratios. Cell shortening was measured optically with a R12 dual raster line edge detector system (Crescent Electronics, Sandy, UT, USA). Images of contracting myocytes were viewed with a charge-coupled device camera mounted with a dual C port adaptor (PTI, USA) on the side port of the microscope. The camera signals were linked to the edge detector electronics. All signals were collected at a sampling rate of 200 points/s.

### Whole-cell patch-clamp recording

A small aliquot of dissociated cells were placed in a 1-mL chamber mounted on the stage of an inverted microscope (Axio Observer Z1, Carl Zeiss). Cells were bathed at room temperature (25–27°C) in normal Tyrode solution. Ionic currents were recorded in whole-cell configuration as described previously [16]. Patch electrodes were made from glass capillaries (o.d.: 1.5 mm, i.d.: 1.0 mm; A-M Systems, Sequim, WA, USA) using a 2-stage vertical puller (P-830, Narishige, Tokyo, Japan) and were fire polished. The resistances of the electrode were 2–5 MΩ when filled with normal pipette solution (containing in μM: KCl 120.0, NaCl 10.0, MgATP 5.0, EGTA 5.0, and HEPES 10.0, adjusted to pH 7.2 with KOH). Membrane currents were recorded using a voltage clamp amplifier (Axopatch 200B, Molecular Devices, Sunnyvale, CA, USA). Electrode junction potentials (5–10 mV) were measured and nulled before suction of the cell. A high-resistance seal (5–10 GΩ) was obtained before the disruption of the membrane patch. Usually, more than 5 min was allowed for adequate cell dialysis following disruption of the membrane patch and before initiating the voltage pulse protocol. Series resistances were compensated to minimize the duration of capacitive surge on the current recording and the voltage drop produced across the clamped cell membrane. About 60–80% of series resistances were compensated. Cell capacitance was measured by calculating the total charge movement of the capacitive transient in response to a 5-mV hyperpolarizing pulse. Command pulses were generated by a 12-bit digital-to-analog converter (Digidata 1320A, Molecular Devices) controlled by pCLAMP software (Molecular Devices, version 8).

During measurement of K^+^ currents, Ca^2+^ and Na^+^ inward currents were blocked by the addition of 1 μM Co^2+^ and 30 μM tetrodotoxin (TTX) to the bathing solution, respectively. To activate K^+^ currents, cells were voltage clamped at a potential of −80 mV and currents (inward rectifier [*I*_K1_] or transient outward K^+^ current [*I*_to_]) were elicited by 500-ms hyperpolarizing or depolarizing test pulses ranging from −140 to +60 mV. Steady-state inactivation of *I*_to_ was examined with a double-pulse protocol: a conditioning 400-ms pulse to various potentials ranging from −80 to 0 mV was followed by a test depolarizing pulse to +60 mV. The holding potential was −80 mV. The twin-pulse protocol, which consisted of 2 identical 200-ms depolarizing pulses to +60 mV from a holding potential of −80 mV, was used to study the recovery of *I*_to_ channels. Prepulse–test pulse intervals varied between 10 and 550 ms.

### Statistical methods and data analysis

Continuous data are presented as mean ± standard deviation (S.D.) unless otherwise indicated. Categorical data are expressed as frequency (%). Statistical comparisons were made using SAS 9.1 (SAS Institute Inc., Cary, NC, USA) and *p* values < 0.05 were considered statistically significant. Student’s *t*-test was used for univariate analysis of continuous variables. Generalized estimation equation (GEE) models were conducted to analyze the change of outcome variables over time. The body weight of rats was also adjusted in fitting GEE models. The distribution of outcome variables was proved to be Gaussian by a normality test. Exchangeable working correlation matrix was applied when applying the GEE method. Concentration–response curves were fitted by an equation of the form: *E*= *E*_max_/[1+(*IC*_50_)^*n*H^], where *E* is the effect at concentration C, *E*_max_ is maximal effect, *IC*_50_ is the concentration for half-maximal inhibition and *n*_H_ is the Hill coefficient. Conductance of *I*_to_ (*G*_to_) was calculated according to the equation as follows: *G*_to_ = *I*_to_/(*V*_m_–*V*_rev_), where *V*_rev_ is the reversal potential of *I*_to_. The activation curves of *I*_to_ were fitted by the Boltzmann equation: *G*_to_/*G*_to, max_ = 1/{1+exp[(*V*_h_-*V*_m_)/*k*]}, where *G*_to, max_ is the maximal ionic conduction, *V*_h_ is the half-maximal activation potential, *V*_m_ is the membrane potential, and k is the slope factor. The inactivation curves of *I*_to_ were fitted by the Boltzmann equation: *I*_to_/*I*_to, max_ = 1/{1+exp[(*V*_m_-*V*_h_)/*k*]}; where *I*_to_ gives the current amplitude and *I*_to, max_ its maximum, *V*_m_ the potential of prepulse, *V*_h_ the half-maximal inactivation potential, and k the slope factor.

## Results

### Effect of PQ on hemodynamic measurements and the ECG

Baseline hemodynamic and ECG parameters did not vary significantly among rats receiving the vehicle or PQ at a dose of 100 and 180 mg/kg (Table S1). Figure 1 is a representative sample of the effect of PQ (180 mg/kg) on arterial pressure, LV pressure, first derivative of LV pressure (LV d*P*/d*t*), and the ECG at different time points. PQ impeded left ventricle performance in both systolic and diastolic phases, decreased blood pressure, slow heart rate, prolonged QT and QTc intervals in a dose-dependent manner. (Table 1)

**Table 1.**
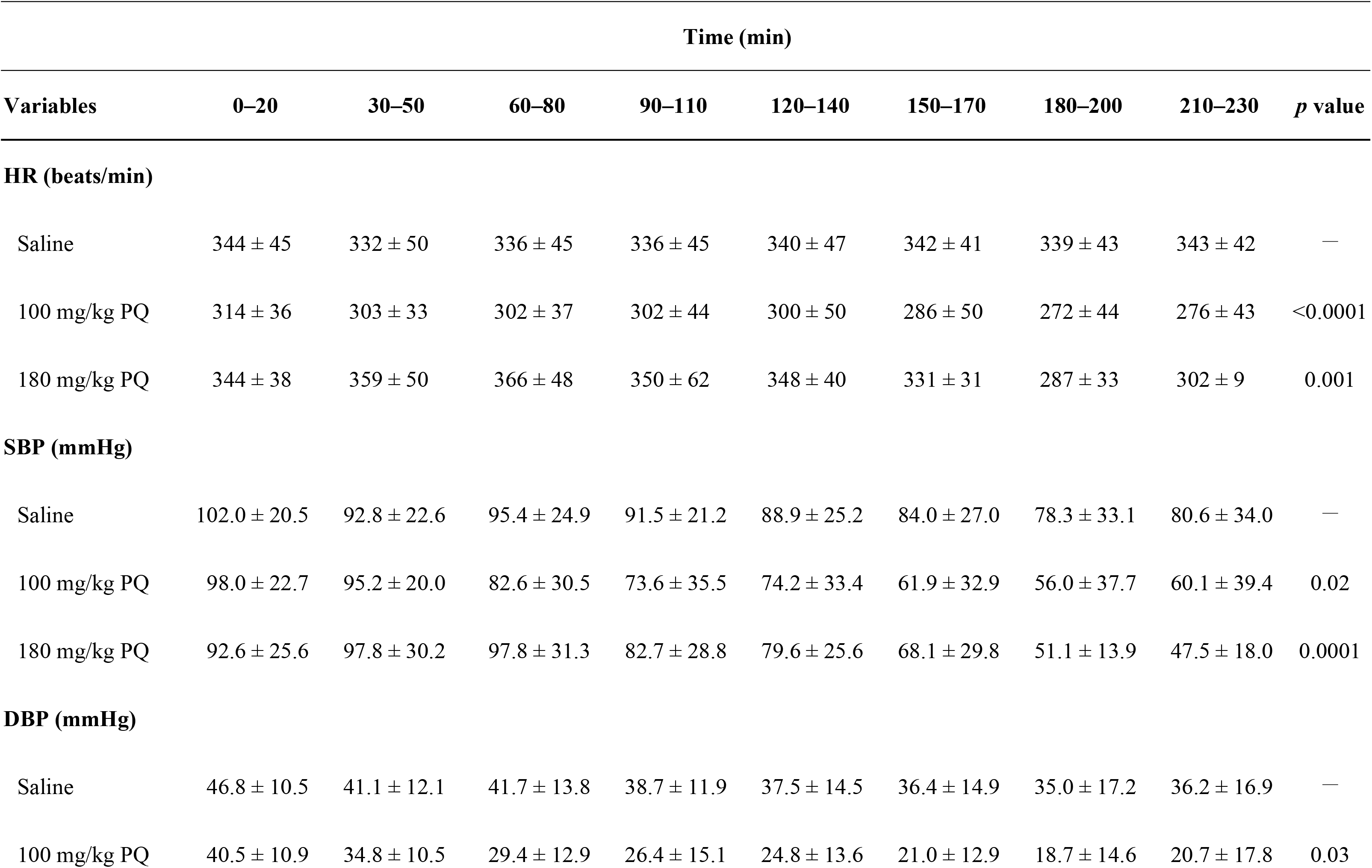
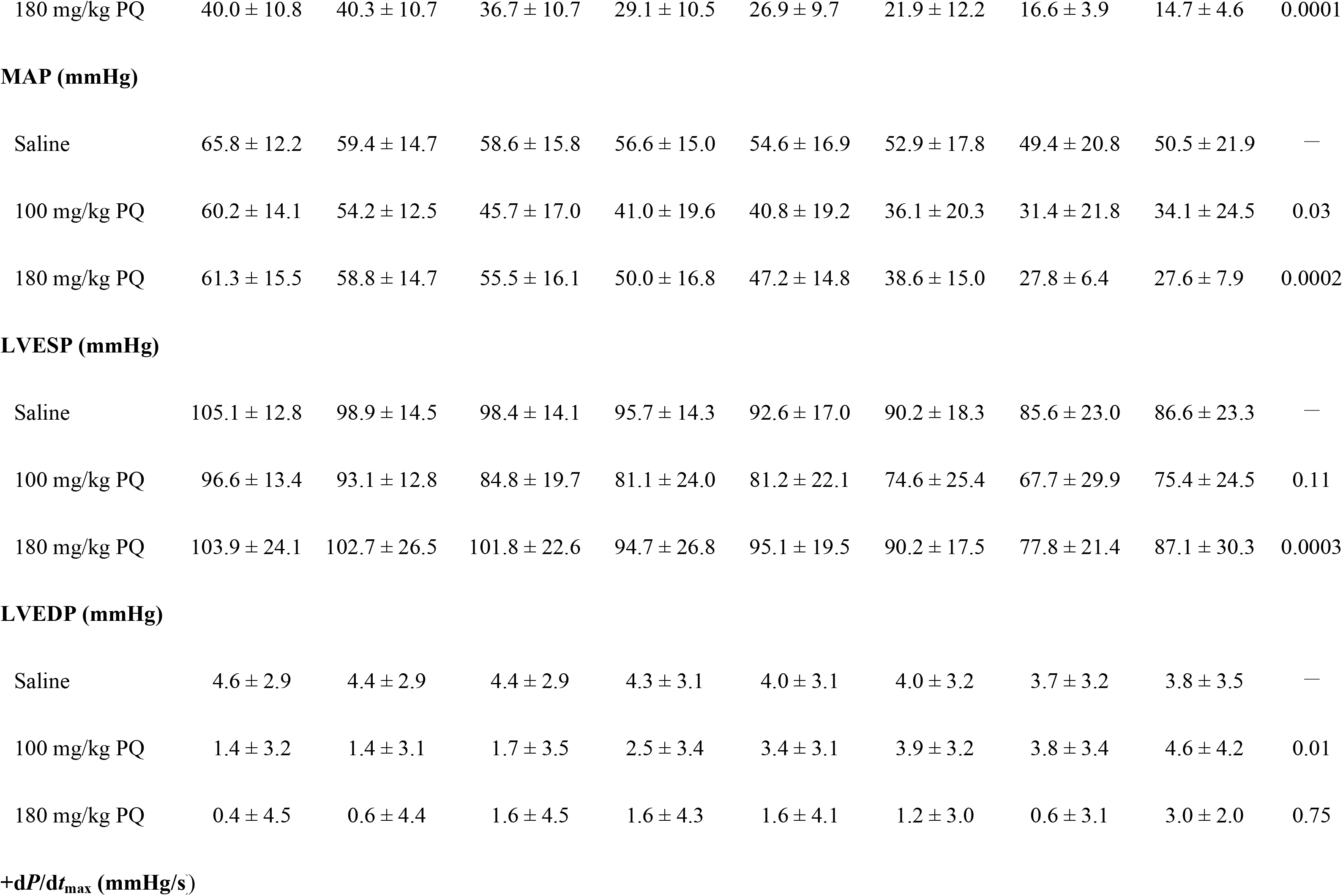
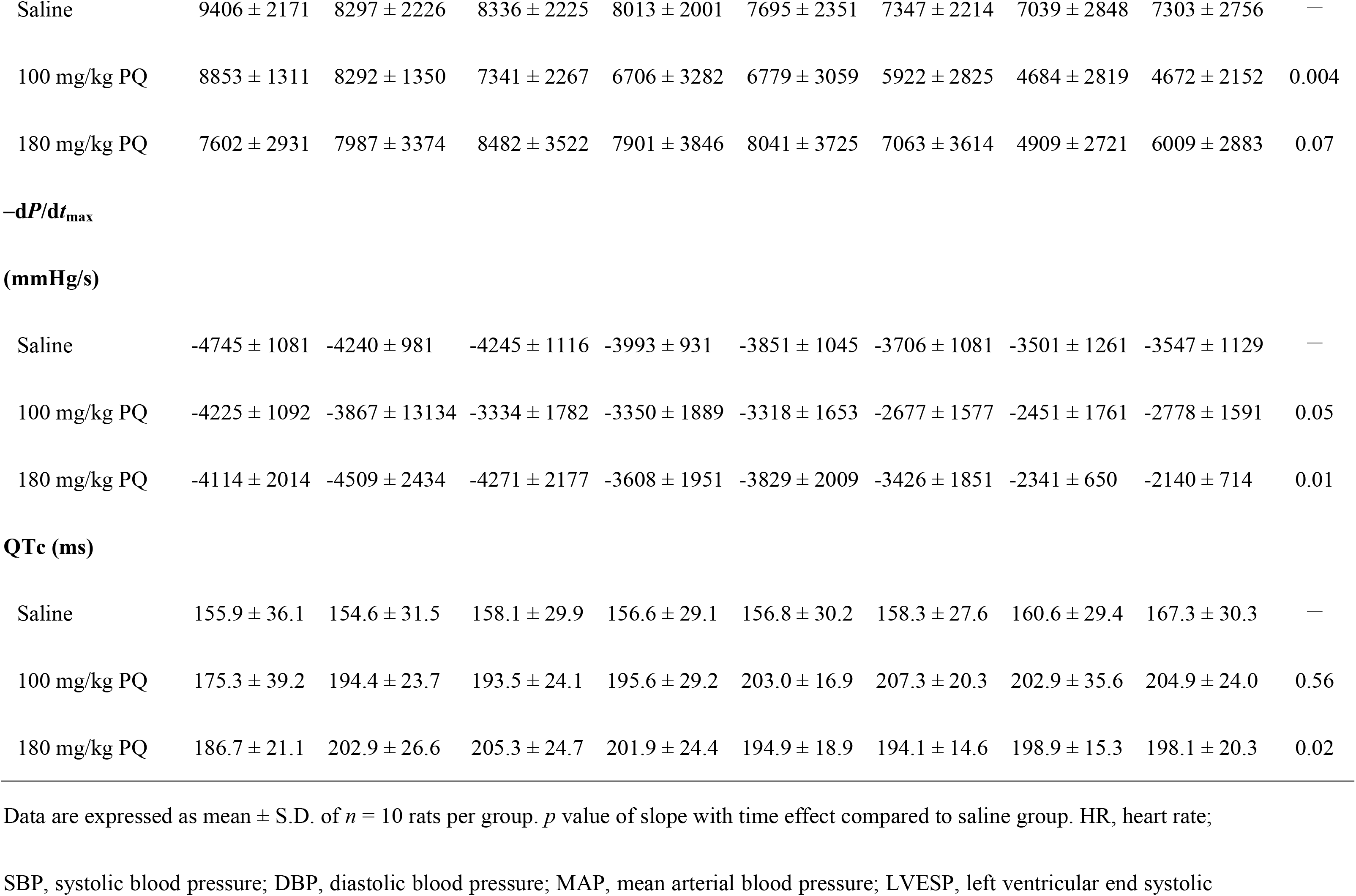
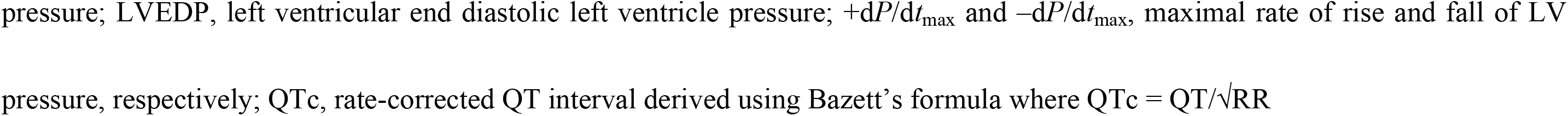
Effects of PQ on hemodynamic and electrocardiographic variables in anesthetized rats.

|  | Time (min) |  |  |  |  |  |  |  |  |
| --- | --- | --- | --- | --- | --- | --- | --- | --- | --- |
| Variables | 0–20 | 30–50 | 60–80 | 90–110 | 120–140 | 150–170 | 180–200 | 210–230 | *p* value |
| HR (beats/min) |  |  |  |  |  |  |  |  |  |
| Saline | 344 ± 45 | 332 ± 50 | 336 ± 45 | 336 ± 45 | 340 ± 47 | 342 ± 41 | 339 ± 43 | 343 ± 42 | — |
| 100 mg/kg PQ | 314 ± 36 | 303 ± 33 | 302 ± 37 | 302 ± 44 | 300 ± 50 | 286 ± 50 | 272 ± 44 | 276 ± 43 | <0.0001 |
| 180 mg/kg PQ | 344 ± 38 | 359 ± 50 | 366 ± 48 | 350 ± 62 | 348 ± 40 | 331 ± 31 | 287 ± 33 | 302 ± 9 | 0.001 |
| SBP (mmHg) |  |  |  |  |  |  |  |  |  |
| Saline | 102.0 ± 20.5 | 92.8 ± 22.6 | 95.4 ± 24.9 | 91.5 ± 21.2 | 88.9 ± 25.2 | 84.0 ± 27.0 | 78.3 ± 33.1 | 80.6 ± 34.0 | — |
| 100 mg/kg PQ | 98.0 ± 22.7 | 95.2 ± 20.0 | 82.6 ± 30.5 | 73.6 ± 35.5 | 74.2 ± 33.4 | 61.9 ± 32.9 | 56.0 ± 37.7 | 60.1 ± 39.4 | 0.02 |
| 180 mg/kg PQ | 92.6 ± 25.6 | 97.8 ± 30.2 | 97.8 ± 31.3 | 82.7 ± 28.8 | 79.6 ± 25.6 | 68.1 ± 29.8 | 51.1 ± 13.9 | 47.5 ± 18.0 | 0.0001 |
| DBP (mmHg) |  |  |  |  |  |  |  |  |  |
| Saline | 46.8 ± 10.5 | 41.1 ± 12.1 | 41.7 ± 13.8 | 38.7 ± 11.9 | 37.5 ± 14.5 | 36.4 ± 14.9 | 35.0 ± 17.2 | 36.2 ± 16.9 | — |
| 100 mg/kg PQ | 40.5 ± 10.9 | 34.8 ± 10.5 | 29.4 ± 12.9 | 26.4 ± 15.1 | 24.8 ± 13.6 | 21.0 ± 12.9 | 18.7 ± 14.6 | 20.7 ± 17.8 | 0.03 |
| 180 mg/kg PQ | 40.0 ± 10.8 | 40.3 ± 10.7 | 36.7 ± 10.7 | 29.1 ± 10.5 | 26.9 ± 9.7 | 21.9 ± 12.2 | 16.6 ± 3.9 | 14.7 ± 4.6 | 0.0001 |
| **MAP (mmHg)** |  |  |  |  |  |  |  |  |  |
| Saline | 65.8 ± 12.2 | 59.4 ± 14.7 | 58.6 ± 15.8 | 56.6 ± 15.0 | 54.6 ± 16.9 | 52.9 ± 17.8 | 49.4 ± 20.8 | 50.5 ± 21.9 | — |
| 100 mg/kg PQ | 60.2 ± 14.1 | 54.2 ± 12.5 | 45.7 ± 17.0 | 41.0 ± 19.6 | 40.8 ± 19.2 | 36.1 ± 20.3 | 31.4 ± 21.8 | 34.1 ± 24.5 | 0.03 |
| 180 mg/kg PQ | 61.3 ± 15.5 | 58.8 ± 14.7 | 55.5 ± 16.1 | 50.0 ± 16.8 | 47.2 ± 14.8 | 38.6 ± 15.0 | 27.8 ± 6.4 | 27.6 ± 7.9 | 0.0002 |
| **LVESP (mmHg)** |  |  |  |  |  |  |  |  |  |
| Saline | 105.1 ± 12.8 | 98.9 ± 14.5 | 98.4 ± 14.1 | 95.7 ± 14.3 | 92.6 ± 17.0 | 90.2 ± 18.3 | 85.6 ± 23.0 | 86.6 ± 23.3 | — |
| 100 mg/kg PQ | 96.6 ± 13.4 | 93.1 ± 12.8 | 84.8 ± 19.7 | 81.1 ± 24.0 | 81.2 ± 22.1 | 74.6 ± 25.4 | 67.7 ± 29.9 | 75.4 ± 24.5 | 0.11 |
| 180 mg/kg PQ | 103.9 ± 24.1 | 102.7 ± 26.5 | 101.8 ± 22.6 | 94.7 ± 26.8 | 95.1 ± 19.5 | 90.2 ± 17.5 | 77.8 ± 21.4 | 87.1 ± 30.3 | 0.0003 |
| **LVEDP (mmHg)** |  |  |  |  |  |  |  |  |  |
| Saline | 4.6 ± 2.9 | 4.4 ± 2.9 | 4.4 ± 2.9 | 4.3 ± 3.1 | 4.0 ± 3.1 | 4.0 ± 3.2 | 3.7 ± 3.2 | 3.8 ± 3.5 | — |
| 100 mg/kg PQ | 1.4 ± 3.2 | 1.4 ± 3.1 | 1.7 ± 3.5 | 2.5 ± 3.4 | 3.4 ± 3.1 | 3.9 ± 3.2 | 3.8 ± 3.4 | 4.6 ± 4.2 | 0.01 |
| 180 mg/kg PQ | 0.4 ± 4.5 | 0.6 ± 4.4 | 1.6 ± 4.5 | 1.6 ± 4.3 | 1.6 ± 4.1 | 1.2 ± 3.0 | 0.6 ± 3.1 | 3.0 ± 2.0 | 0.75 |
| **+dP/dt<sub>max</sub> (mmHg/s)** |  |  |  |  |  |  |  |  |  |
| Saline | 9406 ± 2171 | 8297 ± 2226 | 8336 ± 2225 | 8013 ± 2001 | 7695 ± 2351 | 7347 ± 2214 | 7039 ± 2848 | 7303 ± 2756 | — |
| 100 mg/kg PQ | 8853 ± 1311 | 8292 ± 1350 | 7341 ± 2267 | 6706 ± 3282 | 6779 ± 3059 | 5922 ± 2825 | 4684 ± 2819 | 4672 ± 2152 | 0.004 |
| 180 mg/kg PQ | 7602 ± 2931 | 7987 ± 3374 | 8482 ± 3522 | 7901 ± 3846 | 8041 ± 3725 | 7063 ± 3614 | 4909 ± 2721 | 6009 ± 2883 | 0.07 |
| **−dP/dt<sub>max</sub>** |  |  |  |  |  |  |  |  |  |
| **(mmHg/s)** |  |  |  |  |  |  |  |  |  |
| Saline | −4745 ± 1081 | −4240 ± 981 | −4245 ± 1116 | −3993 ± 931 | −3851 ± 1045 | −3706 ± 1081 | −3501 ± 1261 | −3547 ± 1129 | — |
| 100 mg/kg PQ | −4225 ± 1092 | −3867 ± 13134 | −3334 ± 1782 | −3350 ± 1889 | −3318 ± 1653 | −2677 ± 1577 | −2451 ± 1761 | −2778 ± 1591 | 0.05 |
| 180 mg/kg PQ | −4114 ± 2014 | −4509 ± 2434 | −4271 ± 2177 | −3608 ± 1951 | −3829 ± 2009 | −3426 ± 1851 | −2341 ± 650 | −2140 ± 714 | 0.01 |
| **QTc (ms)** |  |  |  |  |  |  |  |  |  |
| Saline | 155.9 ± 36.1 | 154.6 ± 31.5 | 158.1 ± 29.9 | 156.6 ± 29.1 | 156.8 ± 30.2 | 158.3 ± 27.6 | 160.6 ± 29.4 | 167.3 ± 30.3 | — |
| 100 mg/kg PQ | 175.3 ± 39.2 | 194.4 ± 23.7 | 193.5 ± 24.1 | 195.6 ± 29.2 | 203.0 ± 16.9 | 207.3 ± 20.3 | 202.9 ± 35.6 | 204.9 ± 24.0 | 0.56 |
| 180 mg/kg PQ | 186.7 ± 21.1 | 202.9 ± 26.6 | 205.3 ± 24.7 | 201.9 ± 24.4 | 194.9 ± 18.9 | 194.1 ± 14.6 | 198.9 ± 15.3 | 198.1 ± 20.3 | 0.02 |

Data are expressed as mean ± S.D. of $n = 10$ rats per group. $p$ value of slope with time effect compared to saline group. HR, heart rate;
SBP, systolic blood pressure; DBP, diastolic blood pressure; MAP, mean arterial blood pressure; LVESP, left ventricular end systolic
pressure; LVEDP, left ventricular end diastolic left ventricle pressure; $+dP/dt_{\max}$ and $-dP/dt_{\max}$ , maximal rate of rise and fall of LV
pressure, respectively; QTc, rate-corrected QT interval derived using Bazett's formula where $QTc = QT/\sqrt{RR}$

**Figure 1.**
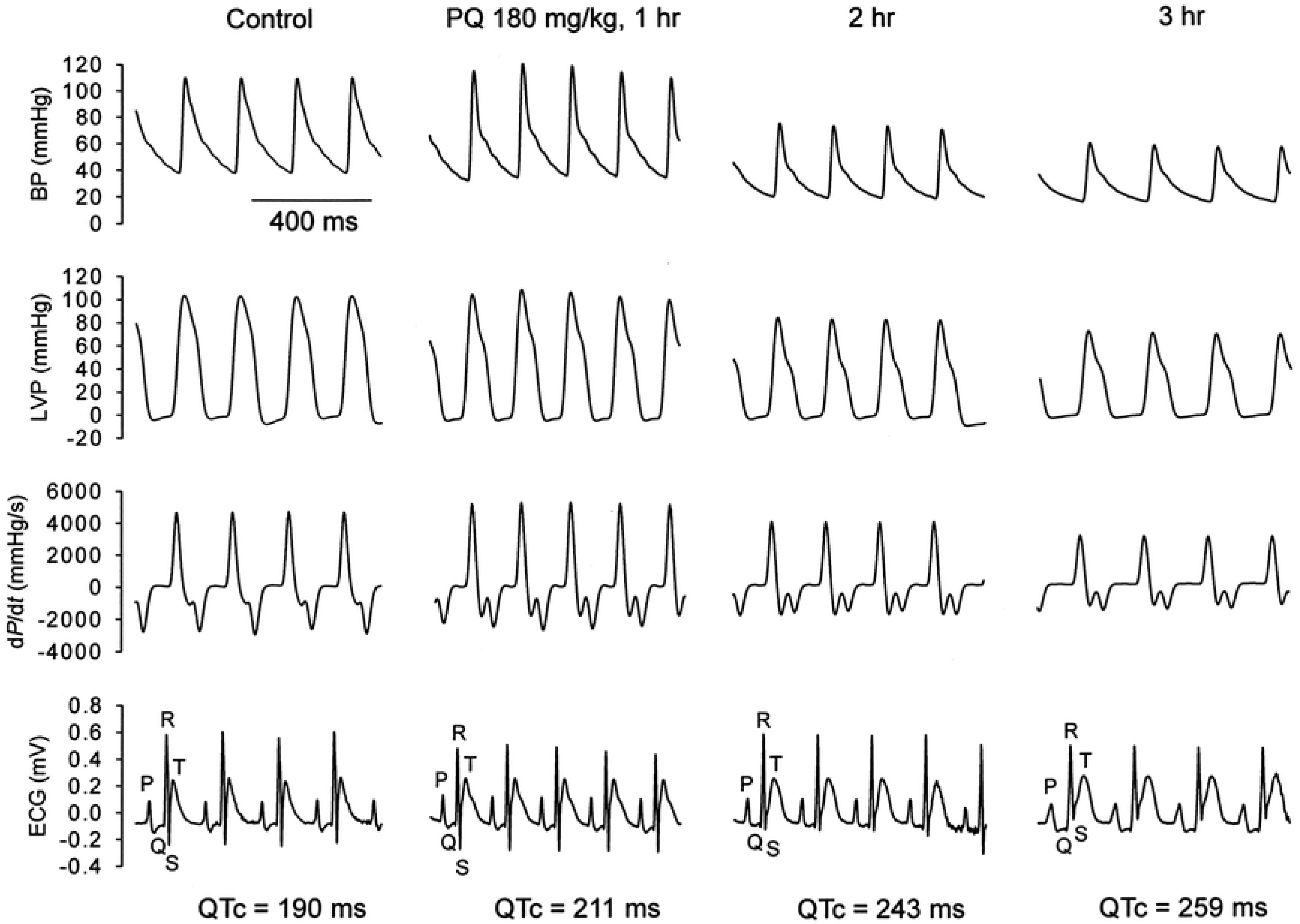
Representative recordings of arterial pressure, LV pressure (LVP), first derivative of LV pressure (LV d*P*/d*t*), and ECG from an anesthetized rat at baseline and at various time points after treatment with PQ (180 mg/kg, i.v.).

### Effect of PQ on contractile force in Langendorff-perfused rat hearts

PQ had negative effects on contractile force in Langendorff-perfused rat hearts. As shown in Figure 2, PQ decreased LV developed pressure (LVDP) and the rate of rise (+d*P*/d*t*_max_,) and fall (−d*P*/d*t*_max_,) of LV pressure in isolated hearts in a concentration-dependent manner. On average (*n* = 10 for both groups), LVDP decreased from a baseline value of 30.1 ± 10.7 to 18.1 ± 6.9 mmHg and from 25.6 ± 6.4 to 8.5 ± 5.1 mmHg after 60 min treatment with 33 and 60 μM PQ, respectively (*p* < 0.0001). LV + d*P*/d*t*_max_, decreased from a baseline value of 950.7 ± 286.3 to 581.6 ± 183.7 mmHg/s and from 773.3 ± 164.8 to 272.4 ± 167.7 mmHg/s with the application of 33 and 60 μM PQ, respectively (*p* < 0.0001). LV −d*P*/d*t*_max_, decreased from 600.8 ± 271.4 to 387.1 ± 167.5 mmHg/s and from 532.1 ± 116.4 to 177.0 ± 105.4 mmHg/s with the application of 33 and 60 μM PQ, respectively (*p* < 0.0001).

**Figure 2.**
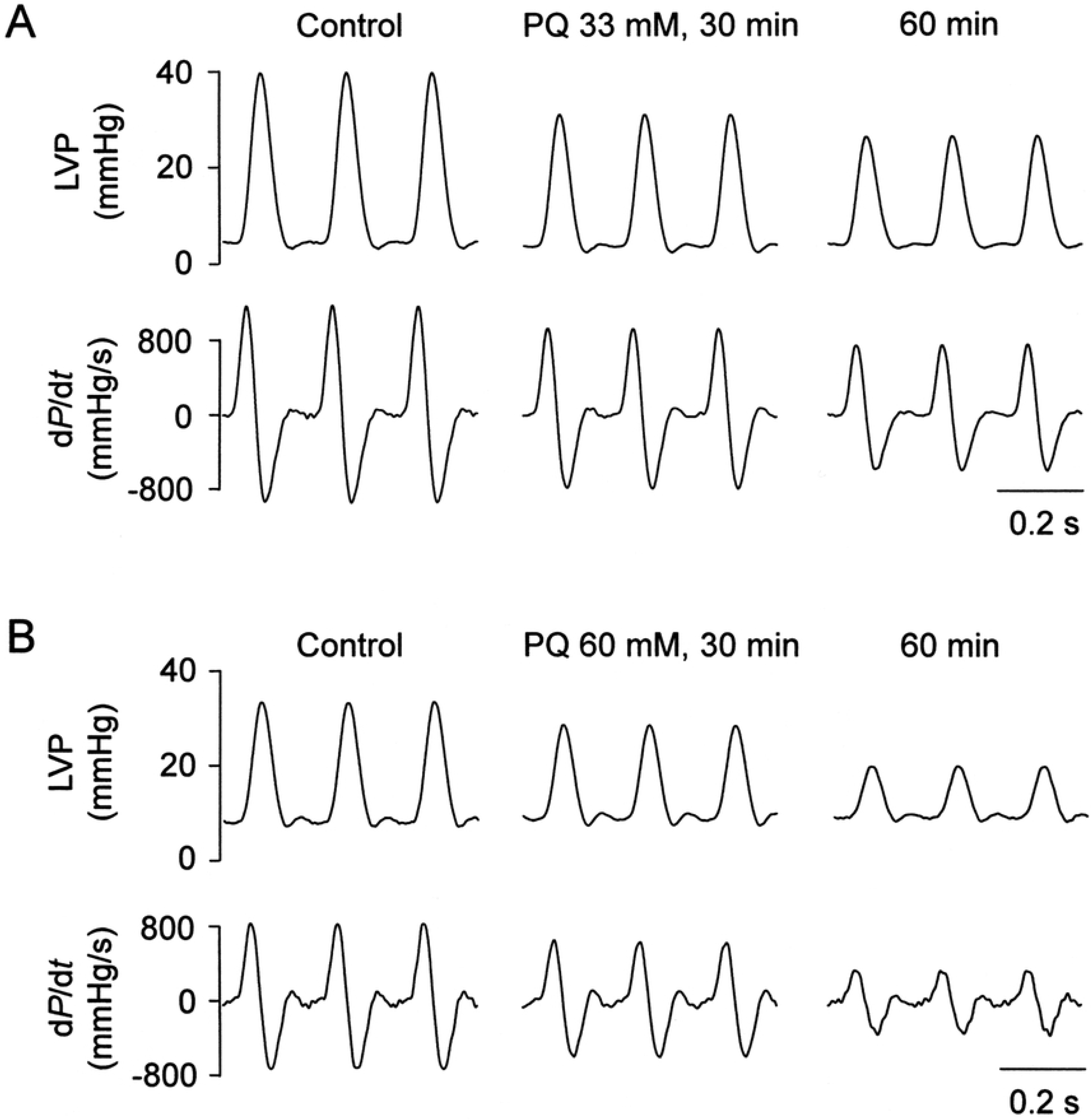
Representative LVP signals recorded from Langendorff-perfused rat hearts at baseline and following treatment with 33 μM (panel A) or 60 μM PQ (panel B).

### Effects of PQ on intracellular Ca^2+^ transients and cell shortening in rat ventricular myocytes

Figure 3A shows the continuous, expanded recordings (Figure 3B), and the summarized data (Figure 3C and 3D) of the effects of PQ (10–60 μM) on Ca^2+^ transients (fura-2 fluorescence ratio *F*_340_/*F*_380_) and cell shortening in rat ventricular myocytes. PQ decreased cell shortening in a concentration-dependent manner (Figure 3C). However, PQ had no effect on time-to-peak of cell shortening and time to 50% relengthening (Figure S1A and S1B). PQ decreased the amplitude of fluorescence ratio (Ca^2+^ transients) in a concentration-dependent manner (Figure 3D). However, it had no obvious effect on time to peak Ca^2+^ transients (Figure S1C). At concentrations of 10 and 60 μM, PQ significantly decreased the decay time constant of Ca^2+^ transients, while it showed no significant effect at 30 μM (Figure S1D).These findings suggested that decreases contractility force of cardiac myocytes but not their peak shortening/relengthening velocities.

**Figure 3.**
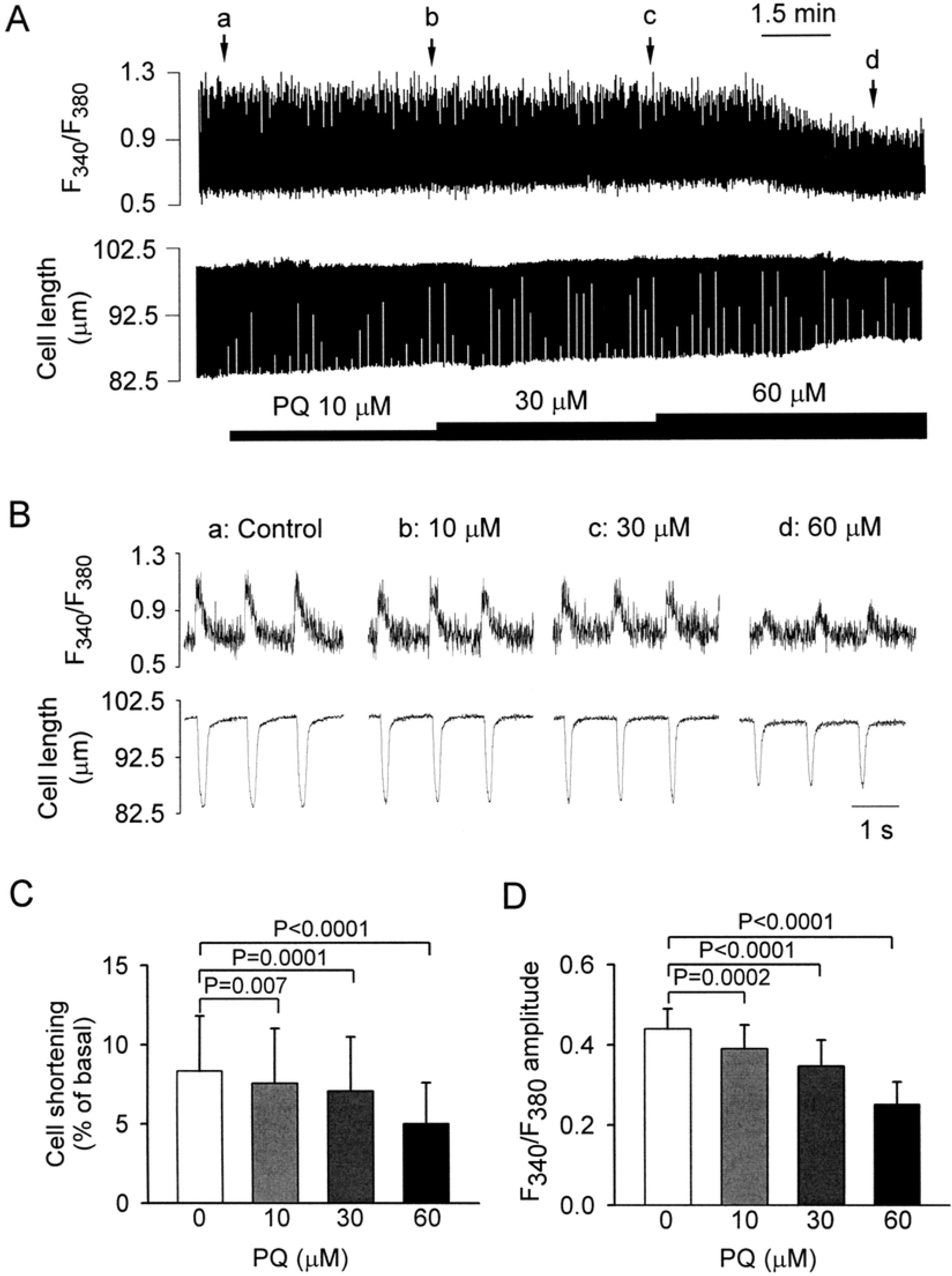
Effect of PQ on Ca^2+^ transients (represented by fura-2 fluorescence ratio *F*_340_/*F*_380_) and cell shortening in rat ventricular myocytes. (A) Continuous recordings of Ca^2+^ transients (upper panel) and cell shortening (lower panel) showing the effects of cumulative application of 10, 30, and 60 μM PQ. (B) Recordings on an expanded time scale taken at the time indicated by the corresponding letters over the *F*_340_/*F*_380_ trace in panel A. (C, D) Mean data of the amplitude of cell shortening (C) and Ca^2+^ transient (D) before and after application of PQ. Data are expressed as mean ± S.D. (*n* = 11). Cell shortening was normalized to resting cell length.

### Effects of PQ on K^+^ currents in rat isolated ventricular myocytes

In order to separate the K^+^ currents from overlapping currents, Na^+^ and Ca^2+^ currents were blocked with 30 μM TTX and 1 μM Co^2+^, respectively. Typical current traces recorded in response to depolarizing and hyperpolarizing clamp steps to test potentials between +60 and −140 mV from a holding potential of −80 mV are shown in Figure 4A. PQ (3 and 10 μM) moderately reduced the amplitude of peak outward K^+^ current (*I*_to_) but had no significant effect on inward rectifier K^+^ current (*I*_K1_). PQ also reduced the amplitude of steady-state outward K^+^ current (*I*_ss_) at the end of 400-ms long clamp steps. Figure 4B and 4C shows the current density–voltage (*I*−*V*) relationship for the peak currents and *I*_ss_ before and after the addition of PQ (1, 3, and 10 μM), respectively. Figure 4D shows the percent inhibition of *I*_to_ integral by PQ was not significantly dependent on the step potential.

**Figure 4.**
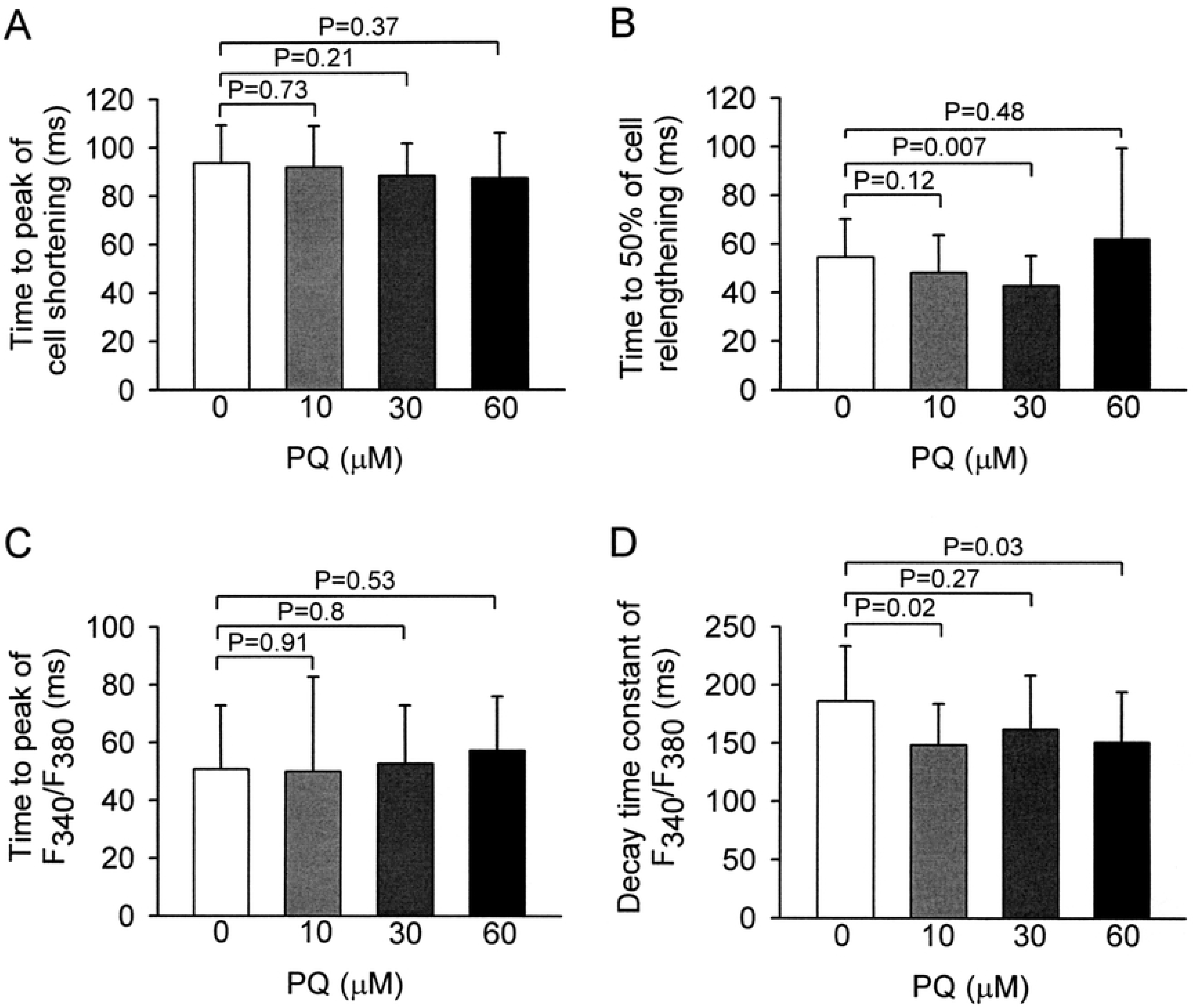
Effect of PQ on K^+^ currents. (A) Families of current traces elicited by a series of 400-ms long depolarizing or hyperpolarizing pulses from a holding potential of −80 mV in the absence and presence of 1, 3, and 10 μM PQ. Arrow in each panel indicates zero current level. (B, C) Averaged I–V relationship for peak currents (B) and *I*_ss_ (C) observed in the absence, and presence of 1, 3, and 10 μM PQ. Each data point indicates mean ± S.D. from 5 myocytes. (D) Percent inhibition of the *I*_to_ integral by 1, 3, and 10 μM PQ calculated at different depolarizing potentials. Data points are mean ± S.D. (*n* = 5). (E) Original **s**uperimposed families of current traces generated by 400-ms depolarizing pulses to +40 mV from a holding potential of −80 mV in the absence or presence of increasing concentrations of PQ. Arrow indicates zero current level. (F) Concentration–response curve for the effect of PQ on the integral of *I*_to_ at +40 mV. Data points are mean ± S.D. (*n* = 5). The continuous line was drawn according to the fitting of Hill equation.

The effect of PQ on *I*_to_ was investigated further by analyzing its concentration dependence. *I*_to_ was elicited by a depolarizing pulse to +40 mV from a holding potential of −80 mV. Figure 4E shows superimposed K^+^ current traces before and after cumulative superfusion with 3 and 10 μM PQ. The decay of currents during an activating clamp step in control conditions and after administration with 1, 3, and 10 μM PQ were well fitted by a single exponential function: *I*_to_ (t) = A_1_ exp (−t/τ) + A_0_, where A_1_ and τ are the initial amplitude and time constant of inactivation, respectively. A_0_ is a time-independent component. Under control conditions, the average value of the decay time constant (τ) was 38.7 ± 13.4 ms (*n* = 5). In the presence of 1, 3, and 10 μM PQ, decay τ was 37.7 ± 1.3, 49.3 ± 16.2, and 52.1 ± 19.8 ms, respectively (*p* = 0.26, *n* = 5). PQ had no significant effect on the decay time course of *I*_to_. Figure 4F illustrates the percent reduction of the *I*_to_ integral as a function of the logarithm of PQ concentration. The data were fitted with a Hill equation to obtain a concentration– response curve. The calculated *IC*_50_ for *I*_to_ was 2.4 μM, with an *E*_max_ of 35.4% inhibition and a *n*_H_ of 2.4 (*n* = 5).

### Effects of PQ on steady-state activation, inactivation, and recovery from inactivation of *I*to

The predrug superimposed current traces are shown in Figure 5A, and the voltage dependence of steady-state activation and inactivation curve of *I*_to_ is shown in Figure 5B. The steady-state inactivation relationship was obtained using a conventional double-pulse protocol. Each peak current was normalized to the maximum current measured and plotted as a function of the conditioning potential. The resultant curves were fitted by the Boltzmann equation to estimate half-inactivation potential (*V*_h_) and slope factor (*k*). As shown in Figure 5B, PQ caused a leftward shift of the steady-state inactivation relationship of *I*_to_. Under control conditions (*n* = 5), mean *V*_h_ was calculated as −24.7 ± 9.8 mV and *k* as −4.9 ± 1.9 mV. Mean *V*_h_ was −39.9 ± 5.6, −48.5 ± 6.6, and −50.5 ± 8.1 mV and k was −4.9 ± 2.0, −4.9 ± 2.0, and −4.0 ± 0.3 mV in the presence of 1, 3, and 10 μM PQ, respectively (*p* = 0.007 for *V*_h_ and *p* = 0.7 for *k*). The activation curves shown in Figure 5B were obtained from the normalized conductance of *I*_to_ channels (*G*_to_/*G*_to, max_) calculated from the *I*_to_ amplitude data obtained in Figure 4A and 4B. Mean half-activation potential (*V*_h_) and k values were 14.9 ± 11.9 mV and 10.4 ± 1.6 mV (*n* = 4), respectively, under control conditions. PQ caused a leftward shift of the voltage dependence for activation to negative potentials. Mean *V*_h_ was −2.5 ± 7.7, −14.4 ± 15.2, and −9.6 ± 9.1 mV and *k* was 11.3 ± 0.9, 18.9 ± 3.9, and 21.4 ± 4.4 mV in the presence of 1, 3, and 10 μM PQ, respectively (*p* = 0.029 for *V*_h_ and p = 0.012 for *k*). The effect of PQ on the recovery kinetics of *I*_to_ was also examined and shown in Figure 5C and 5D. Recovery from inactivation in the control conditions could be well fitted by a single exponential function. As shown in Figure 5D, PQ had no significant effect on the recovery time course of *I*_to_. Mean recovery time constant was 44.2 ± 31.5 ms (*n* = 5) under control conditions. In the presence of 1, 3, and 10 μM PQ, recovery time constants of *I*_to_ were 51.8 ± 33.0, 64.6 ± 45.1, and 94.2 ± 108.3 ms (*p* = 0.38), respectively.

**Figure 5.**
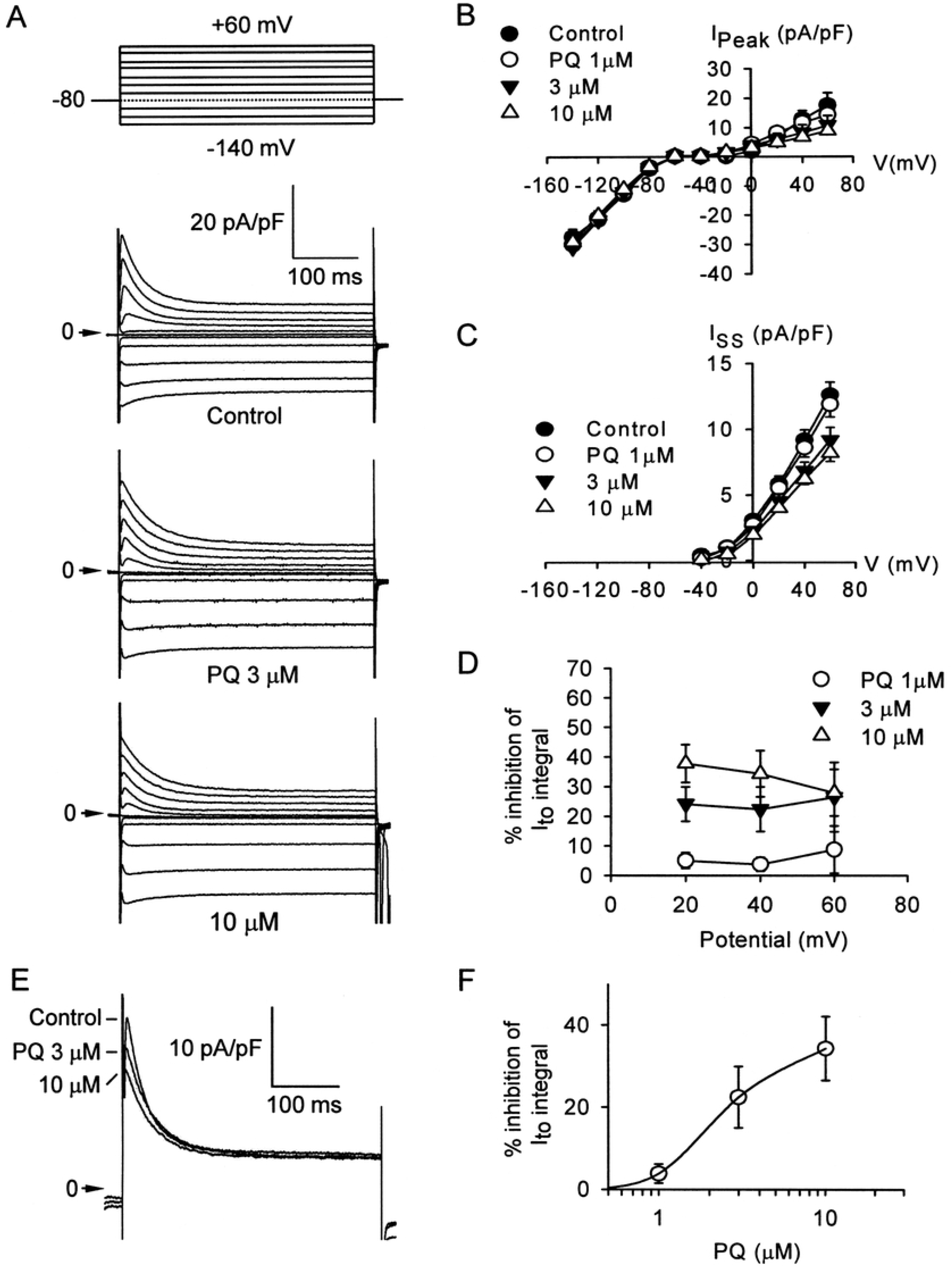
Voltage dependence of steady-state *I*_to_ activation and inactivation in the absence and presence of 3 μM PQ. Steady-state inactivation was examined with a double-pulse protocol: a conditioning 400 ms pulse to various potentials ranging from −80 to +10 mV was followed by a test depolarizing pulse to +60 mV. The holding potential was −80 mV. The predrug superimposed current traces are shown in panel A. The inactivation curves for *I*_to_ were obtained by normalizing the current amplitudes (*I*) to the maximal value (*I*_max_) and plotted as a function of the conditioning potentials before and after PQ (*n* = 5). Solid line drawn through the data points were the best fit to the Boltzmann equation. The activation curves were obtained from the normalized conductance of *I*_to_ channels (*G*_to_/*G*_to, max_), which were calculated from the *I*_to_ amplitude in Figure 4B and plotted as a function of the depolarizing potentials (*n* = 5). The solid line drawn through the data points were the best fit to the Boltzmann equation. Only control and 3 μM PQ were plotted and shown in panel B. (C, D) Effects of PQ on reactivation of *I*_to_. The twin-pulse protocol used consisted of 2 identical 200 ms depolarizing pulses to +60 mV from a holding potential of −80 mV (panel D, inset) and the prepulse–test pulse interval varied between 10 and 550 ms. An example of recovery of *I*_to_ from inactivation in control conditions is shown in panel C. The normalized currents (fractional recovery) obtained in the absence and presence of 3 μM PQ were plotted as a function of the recovery time. Solid lines here represent a single exponential fitted to the data in the absence and presence of 3 μM PQ (*n* = 5), respectively.

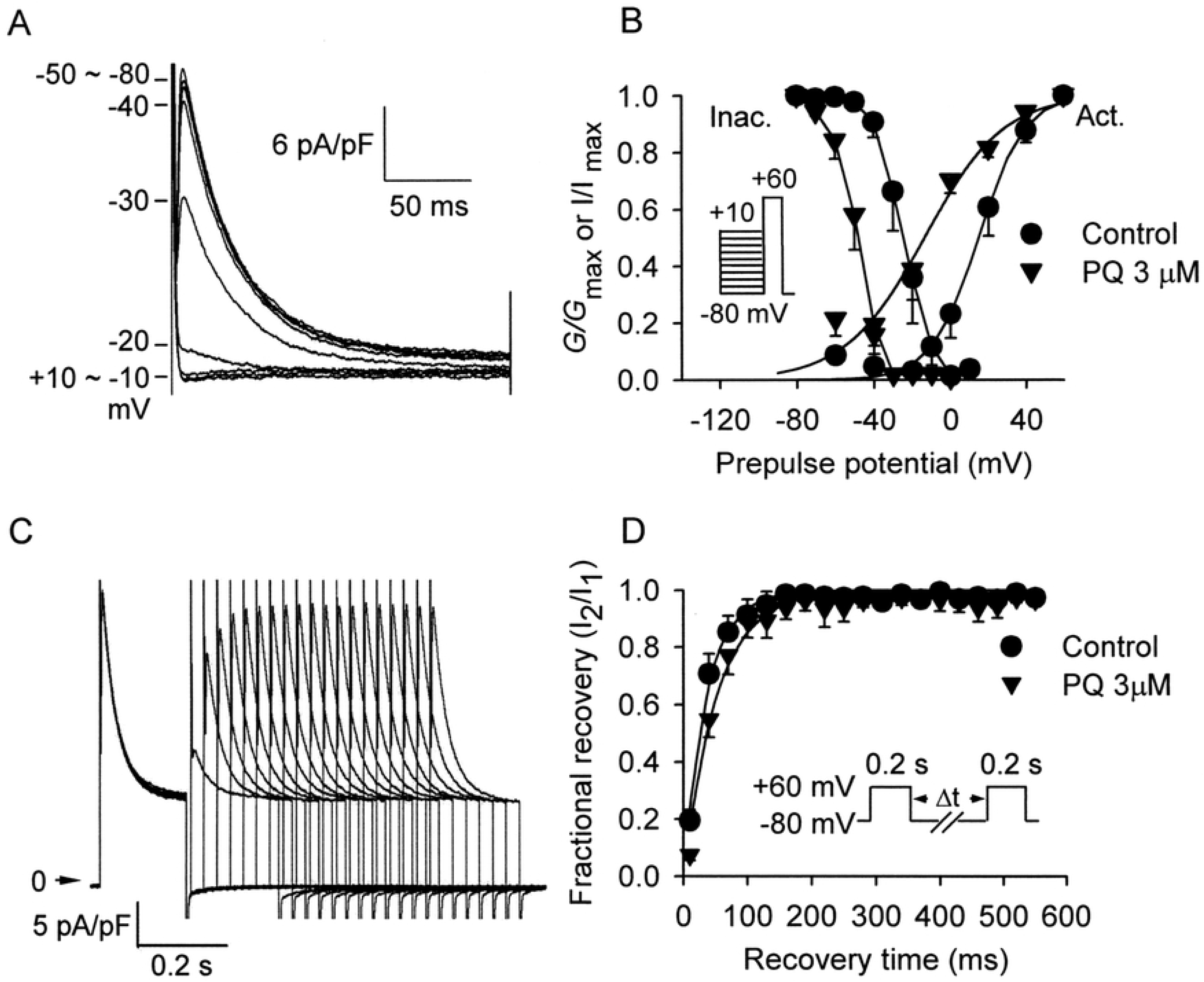

## Discussion

In the present study, hemodynamic and electromechanical effects of paraquat in rat heart were investigated. The major findings of this study are as follows: (1) in anesthetized rats, PQ decreased HR, blood pressure, cardiac contractility, and prolonged the QTc interval in a dose-dependent manner; (2) in Langendorff-perfused rat hearts, PQ decreased LV developed pressure and contractility; (3) in rat ventricular myocytes, PQ decreased both the amplitude of Ca^2+^ transients and cell shortening in a concentration-dependent manner. (4) PQ suppressed *I*_to_ and *I*_ss_ channels. Furthermore, PQ decreased the availability and altered the gating kinetics of *I*_to_ channels. We provide a wide spectrum of PQ effects on heart in this study. PQ not only decrease cardiac contractility but influencing the electrophysiology of heart also.

Clinically, a wide spectrum of cardiovascular effects ranged from minimal changes on the ECG to acute and extensive myocardial necrosis due to acute PQ poisoning has been observed in human PQ poisoning [13]. Severely poisoned patients die from rapid progression of myocardial depression and irreversible circulatory shock in the acute and subacute phases of PQ poisoning [17]. Our study demonstrated that PQ decreased SBP, DBP, MAP, HR, LVESP, and −d*P*/d*t*_max_, and prolonged QT and QTc intervals in a dose-dependent manner in vivo. Furthermore, in vitro, in isolated hearts, PQ decreased LV developed pressure (LVDP) and the rate of rise (+d*P*/d*t*_max_,) and fall (−d*P*/d*t*_max_,) of LV pressure in a concentration-dependent manner, too. PQ also decreased cell shortening in a concentration-dependent manner. The dose-dependent response of PQ observed in our study can be applied to PQ poisoning cases. A rapid accumulation of PQ into the heart but not in the lung or kidney is an important cause of death in the early stage of PQ poisoning when a large amount of paraquat (364 mg/kg) was orally ingested in rats. [10] In human subjects, studies have reported that the dose of PQ is a key prognostic factor for predicting mortality in PQ poisoning patients. [18,19] Patients who suffered from fulminate PQ poisoning mostly within 24–72 h with cardiovascular collapse and multiple organ failure. [6,7]

It is well known that during cardiac excitation–contraction coupling, Ca^2+^ enters the myocyte mainly through L-type Ca^2+^ channels following an action potential and triggers further Ca^2+^ release from the sarcoplasmic reticulum [20]. In the present study, the PQ-induced negative inotropic effect could be explained by its reduction of the amplitude of intracellular Ca^2+^ transients. Depression of contractility was also observed in ventricular myocytes obtained from rats chronically treated with lower doses of PQ [21]. Although the authors observed a reduction in amplitude of cell shortening, they did not observe a change in the amplitude of Ca^2+^ transients. The authors speculated that in using lower PQ doses at chronic exposure, decreased cell shortening could be explained by altered myofilament sensitivity to Ca^2+^. However, in acute- and high-dose PQ exposure, decreased cell shortening could be attributed to the decreased amplitude of Ca^2+^ transients as shown in the present study. In addition to changes in Ca^2+^ transients, PQ-related oxidative stress may cause defects in Ca^2+^ pump and channel functions[22,23], which may also contribute to PQ-induced cardiac dysfunction.

Voltage-gated K^+^ channels play a crucial role in determining the shape and duration of the cardiac action potential. *I*_to_ is generally considered an important repolarizing current in the mammalian action potential, including in human and rat atrium and ventricle [24,25]. Our data showed that *I*_to_ is responsible for the early phase of repolarization and *I*_ss_ which may contribute to the late phase of the rat action potential were differentially blocked by PQ. Kinetic analysis showed that PQ caused a leftward shift of the *I*_to_ inactivation curve and affected the voltage dependence for activation. This finding suggests that PQ may partly bind inactivated channels, which could decrease the number of resting *I*_to_ channels available for activation. The suppression of K^+^ currents could prolong the action potential duration and QTc interval.

Clinically, prolonged QTc interval has been associated with mortality from intoxication with several kinds of pesticides, such as PQ and organophosphates [11, 21,26]. Prolonged QTc interval also affects mortality rates in patients with a variety of cardiac diseases, such as coronary artery disease and congestive heart failure [27,28]. Prolonged QTc interval seems to correlate with poorer LV dysfunction. For example, in patients with anterior acute myocardial infarction, QTc interval might represent an additional marker of LV systolic dysfunction [29]. In patients who received anthracycline treatment, prolonged QTc interval appeared to correlate with LV dysfunction observed by echocardiography [30]. In this context, the observations in our study such as prolongation of the QTc interval and suppression of contractility in isolated hearts, together with decreased cell shortening in ventricular myocytes may contribute to the cardiotoxicity of acute PQ poisoning.

Treating the most severely PQ-intoxicated patients still remains a tremendous challenge in these days. One of the current treatment modalities is immunosuppressive therapy. Some authors claimed that immunosuppressive therapy may have beneficial in treating PQ poisoning patients who died from lung fibrosis-related hypoxemia. [31, 32] However, the methodological problems limit their applicability [33]. Besides, the largest randomized controlled trial completed to date also reported no benefit [34]. We believed that immunosuppressive therapy should have no beneficial in treating the most severe form of PQ poisoned patients which suffered from cardiovascular collapse in the early stage of poisoning. The mechanisms of PQ related dose response cardiac toxicity demonstrated in our study may give us a new view of sight when considering development of new treatment modalities in severe PQ poisoning patients.

## Conclusions

Paraquat has wide negative hemodynamic and electromechanical effects in rat heart. The dose dependent effect on cardiac myocytes contractility and on voltage-gated K^+^ channels could be applied with caution to explain the inevitable death of fulminate paraquat poisoning cases. It seems no efficient treatment modalities could be used to reverse the harmful effects on heart in paraquat poisoning patients in these days.

## Supporting Information

**Table S1 Baseline hemodynamic and electrocardiographic parameters in anesthetized rats receiving the vehicle or PQ at a dose of 100 or 180 mg/kg.**

**Figure S1 Effects of paraquat on kinetic parameters of cell shortening and intracellular Ca^2+^ transients in rat ventricular myocytes.**

## Acknowledgment

This work was supported by Chang Gung Memorial Hospital, Taiwan (grants CMRPG 381571-3) to CCLin. The funders had no role in study design, data collection and analysis, decision to publish, or preparation of the manuscript.

## Author Contributions

CCL:Funding acquisition; CCL and GJC: Conceptualization; CCL and GJC:Methodology; CCL and GJC:Validation; CCL, KHH and GJC:Formal analysis. CCL and GJC Writing - Original Draft/ Review & Editing; All authors read and approved the final manuscript. GJC:Supervision; CCL and GJC are guarantors of the paper.

